# The influence of the respiratory cycle on reaction times in sensory-cognitive paradigms

**DOI:** 10.1101/2021.05.06.442800

**Authors:** Michelle Johannknecht, Christoph Kayser

## Abstract

Behavioural and electrophysiological studies point to widespread influence of the state of respiration on brain activity and cognitive performance. Still, the prevalence and relevance of such respiratory-behavioural relations in typical sensory-cognitive tasks remain unclear. We here used a battery of six tasks probing sensory detection, discrimination and short-term memory to address the questions of whether and by how much behaviour covaries with the respiratory cycle. Our results show that participants tended to align their respiratory cycle to the experimental paradigm. Furthermore, their reaction times, but not so much their response accuracy, consistently and significantly covaried with the respiratory cycle, and this effect was strongest when analyzed contingent on the respiratory state at participants’ responses. The respective effect sizes where comparable to those seen in many typical neurocognitive experimental manipulations. These results support a prominent relation between respiration and sensory-cognitive function and suggest that sensation is intricately linked to rhythmic bodily or interoceptive functions.

## Introduction

Breathing is a vital and automatic process that can be consciously exploited to adapt to behavioural challenges or to control our bodily and mental state. The brain structures controlling breathing and those sensing the resulting changes in airflow are intricately connected with the limbic (Negro et al. 2018). Thereby, information about the respiratory state is potentially widely available in subcortical and cortical brain regions (Heck et al. 2016). This gives us the potential to consciously adjust breathing such as during singing but may also allow respiratory-related signals to continuously and subconsciously influence perceptual or cognitive functions (Kluger & Gross 2020; Tort et al. 2018;Varga & Heck 2017).

In line with the notion that respiration may influence neocortical function, brain activity in multiple brain regions apparently co-modulates with the respiratory cycle, possibly reflecting the propagation of feedback signals about the respiratory state into sensory-cognitive processes (Perl et al. 2019; Kluger & Gross 2020; Zelano et al. 2016; Herrero et al. 2018). Behavioural studies have shown that animal or human participant’s performance can covary with the respiratory cycle in a variety of tasks, ranging from sensory detection (Flexman et al. 1974; Gallego et al. 1991) to emotion recognition (Arch & Craske 2006; Homma & Masaoka 2008; Vlemincx et al. 2010; Zelano et al. 2016), memory recall (Arshamian et al. 2018; Huijbers et al. 2014; Nakamura et al. 2018; Zelano et al. 2016), or more complex mental tasks (Perl et al. 2019). Also, in motor tasks participants preferentially or more swiftly trigger actions during specific respiratory cycles (Park et al. 2020; Li & Rymer, 2011). Often these effects are stronger for nasal than oral breathing, in line with olfactory structures prominently providing feedback about respiratory action to cortical regions (Zelano, et al., 2016).

While these previous studies collectively suggest that perceptual judgements indeed covary with the state of respiration, the prevalence and relevance of such effects during typical laboratory tasks remains unclear. First, previous studies used diverse methods for registering participants respiratory state (respiratory belts, pneumotachographs, manometers) and different analytical approaches to detect the potential covariation of respiration and behaviour. This makes it difficult to compare results across behavioural assays and to establish effect sizes of the behaviour-respiratory relation. Second, many studies explicitly instructed participants to a specific type of respiration (oral, nasal, deep respiration), which may bias participant’s attention to their own respiration and may amplify potential effects (Varga & Heck 2017; Herrero et al. 2018; Vlemincx et al. 2010). Furthermore, typical laboratory paradigms are often structured around specific event times, such as stimulus onsets or participant’s responses. Previous work has not investigated whether the respiratory state during sensory onsets or around the time of participants’ responses more strongly relates to behavioural performance.

The present study was designed to probe whether and by how much participant’s performance in typical perceptual and memory tasks covaries with the respiratory state. For this we exploited a sensitive measurement of respiratory airflow (Short et al. 2016; Grimaud & Murthy 2018) and quantified the variation of response accuracy and reaction times along the respiratory cycle of human participants in six tasks. Importantly, participants performed these tasks without specific instructions as how to breathe other than breath through their nose as usual. The tasks were either based on previous studies reporting a relation between memory performance or judgments of emotions and respiration (Zelano et al. 2016) or were auditory and visual detection or discrimination tasks employed in our previous work. We analysed the data both when combined across all tasks and for each task individually, probing two main questions: whether participants tend to align their respiratory behaviour to the experimental task, and whether performance (reaction time, response accuracy) systematically covaries with the respiratory cycle.

## Methods

### Participants

A total of 42 adult volunteers participated in this study. The procedures were in accordance with the Declaration of Helsinki and were approved by the ethics committee of Bielefeld University. Participants gave written informed consent and were informed about the general procedures and the nature of the individual tasks prior to participating. Our specific interest to investigate the relation between respiration and task performance was not mentioned explicitly prior to the study. A total of six paradigms was used and the data for these were collected in two independent groups of participants. We initially aimed to obtain at least 20 valid datasets for each paradigm, based on general recommendations for behavioral studies (Simmons et al., 2011). However, this goal was not reached for one paradigm (see below). In practice, we collected data from 21 participants in group 1 (emotion discrimination, visual memory, and sound detection; 12 females, mean age 25.5 ± 3.3 years), and from 21 participants in group 2 (two pitch discrimination and a visual motion task; 13 females, mean age 24.2 ± 3 years). Some individual datasets had to be excluded, as described below. Participants received 10€ per hour as compensation for their time.

### General procedures

The experiments were performed in a darkened and sound-proof booth (E:Box; Desone, Germany). Participants sat comfortably in front of a monitor (27” monitor; ASUS PG279Q, 120 Hz refresh rate, gray background of 16 cd/m^2^) and two speakers were positioned adjacent to the left and right of the monitor. Stimulus presentation was controlled using the Psychophysics Toolbox (Version 3.0.14; http://psychtoolbox.org/) using MATLAB (Version R2017a; The MathWorks, Inc., Natick, MA) and was synchronized to an EEG recording system using TTL pulses. Participants responded using a computer keyboard. We also monitored eye movements using an EyeLink 1000 System (monitoring their right eye at 250Hz). However, this was technically not possible for some of the participants and the eye tracking data were not analyzed for this study.

### Recording of respiratory signals

During the main experiments participants wore clinical oxygen masks from which the respiratory-tube connectors were removed. In the opening of these connecters a temperature-sensitive resistor was inserted (Littelfuse Thermistor No. GT102B1K, Mouser electronics). This thermistor allows recording the continuous temperature changes resulting from the respiration-related airflow at high temporal resolution (Short et al. 2016; Grimaud & Murthy 2018). The continuous voltage drop across the thermistor was amplified using a custom-made circuit and recorded and digitized via the analogue input of an ActiveTwo EEG system (BioSemi BV; Netherlands) at a temporal resolution of 500Hz. Participants were instructed to breath normally through the nose, as if performing the experiments without wearing the mask.

### Behavioral paradigms

The following six behavioral paradigms were used. These were administered in two separate groups: Group 1 performed two pitch discrimination tasks (referred to as Pitch1 and Pitch 2 in the following) and a visual motion task (Motion) in counterbalanced order between participants. Group 2 performed an emotion discrimination task (Emotion), a visual memory task (Memory), and sound detection task (Sound), the order of which was the same across participants to ensure the same delay period for the memory part: first was the encoding session for the memory task, then emotion discrimination, then memory retrieval, and finally the sound detection task. In the following we describe each paradigm in detail. Trials started with a fixation period, which was the same for all paradigms (400-1000ms, uniform) unless stated. Inter-trial intervals were the same for all paradigms (1200-1500ms, uniform). Respiratory data was only collected during the main experiments but not during practice blocks or blocks used to determine psychometric thresholds. For each task, participants were instructed to respond as fast and accurately as possible after stimulus presentation.

### Pitch discrimination tasks

The pitch discrimination tasks involved judgements of the pitch of two brief successive tones, as used in two previous studies (Kayser et al. 2016). During each trial two pure tones (50ms duration, 6ms cosine ramp, 50ms pause in between, 65 dB SPL) were presented and participants had two indicate which of the two (first, or second) had higher pitch. One tone was always a standard (1024Hz) the other tone varied around this pitch in five octave-spaced levels. These levels were multiples of the participant-specific threshold, which was obtained separately ([0, 0.5, 1, 1.5, 2] * threshold). Trials started with a fixation period, following which the stimuli were presented. The paradigm consisted of three parts: a brief training session, a block to estimate participant’s thresholds, and finally the main task. Participant-specific thresholds were obtained using three-interleaved one-up two-down staircases with multiplicative step-sizes (starting at differences of 0.5, 0.1 and 0.02 octaves respectively). The three individual thresholds were averaged to yield the participant-specific threshold. The tasks Pitch1 and Pitch 2 differed in that during Pitch 1 the stimulus was presented following the random fixation period. During Pitch 2, participants had to manually initialize the start of the trial by pressing a key, which was then followed by a shorter fixation period (300 to 600ms uniform). For Pitch 1 participants performed two blocks of 200 trials each (resulting in 80 repeats per pitch difference), for Pitch 2 participants performed one block of 200 trials (resulting in 40 repeats per difference). For both tasks the data from all n=21 participants could be analyzed.

### Visual random-dot motion task

In this task, participants judged the direction of motion (left-or right-wards) of visual random dot displays. Random dot displays lasted 340ms, subtended 10 degrees of visual angle and contained 1100 limited-lifetime dots (0.2° diameter, 8 frames life-time) moving at 5 degrees per second. The coherence of the dots (fraction of dots moving in the same direction) varied across five levels around participant’s individual thresholds ([0.55, 0.77, 1, 1.22, 1.45] * threshold). As for pitch discrimination, this paradigm started with a practice block, following a block used to determine the individual threshold. Finally, two blocks of the main task with 200 trials each were performed (resulting in 80 repeats per level). As for pitch discrimination, thresholds were determined using three interleaved one-up two-down staircases with multiplicative step-sizes (starting at coherence levels of 0.6, 0.2 and 0.04 respectively). Due to technical problems with the recording of respiration signals in individual blocks, the data from 3 participants had to be excluded (n=18).

### Emotion discrimination task

In this paradigm, which was modelled based on Zelano et al. (Zelano et al. 2016), participants had to categorize individual faces as either displaying an angry or disgusted emotional expression. The images (subtending about 8 degrees, presented for 100 ms) were taken from the FACES database (Ebner et al. 2010), from which we selected 400 middle-ages faces across both genders and the two emotions. Participants performed a brief training block and two blocks of the experiment with 200 trials each (resulting in 200 trials per emotion). The data from all n=21 participants could be analyzed.

### Visual memory task

In this paradigm participants hat do remember a set of images showing diverse objects and later had to recognize those shown previously. The paradigm was modelled based on the study by Zelano et al. (Zelano et al. 2016) and the images were taken from an existing database (Moreno-Martínez & Montoro 2012). During the exposure phase, a total of 164 images (500ms presentation time) were shown in random order and participants had no other task than to remember these. During the test phase, participants were presented with those 164 previous images and a set of 164 novel images, in pseudo-random order. Participant’s task was to indicate whether the image was previously seen or new, whereby they could respond already during the stimulus presentation time. The inter-trial intervals for both phases were 2 – 2.5s. We investigated the respiration data only during the test period. The data from all n=21 participants could be analyzed.

### Sound detection task

This task involved the detection of an acoustic target sound (100 ms sine-wave tone, 1024 Hz) on a white noise background. The target could take one of four levels (signal to noise ratios, defined as relative root-mean-square amplitudes of tone and white noise) and could either be present or absent. The white noise had a level of 65 dB SPL. The four SNRs were spaced around participant-specific thresholds ([0.3, 0.6, 1.3, 3] * threshold), which were determined in separate blocks. The paradigm started with a practice block, following a block used to determine the individual threshold. Finally, two blocks of the main task with 240 trials each were performed. Individual thresholds were determined using three interleaved one-up two-down staircases with multiplicative step-sizes (starting at 0.5, 0.1 and 0.02 respectively). Due to technical problems with the recording of respiration signals data from only n=20 participants could be analyzed.

### Data analysis of respiratory data

The respiratory signals were filtered using 3-rd order Butterworth filters (high pass at 0.05 Hz, low pass at 8 Hz) and subsequently resampled at 100Hz using the FieldTrip toolbox (Oostenveld et al. 2011). To determine individual respiratory cycles, we implemented two procedures to detect local peaks reflecting the peak inhalation in these traces and found that they yielded very comparable results. One procedure detected local peaks in a low-pass filtered version of the data (1Hz) that were at least 500ms apart. Another procedure applied the Hilbert transform to the data, and determined local peaks based on the respective phase variable. These provided highly similar results: the number of detected respiratory cycles differed by only 1.1±0.2 % (mean ± across participants, n=122 across datasets). Individual respiratory cycles were determined based on the data in 7s windows around each peak (Fig. 1A), whereby peaks were included only if the z-scored trace exceeded a level of z=0.5 (Zelano et al. 2016). To define the state of respiration for each time point, we proceeded as follows. The inspiration period was defined as the continuous period with positive slope prior to the local peak (whereby interruptions of the positive slope shorter than 350ms were interpolated). The expiration period was defined as the continuous period with negative slope subsequent to the local peak (again interruptions shorter than 350ms were interpolated). This definition can leave a few time points near exhalation offset and inspiration onset as unassigned to either state (see below).

**Figure 1.**
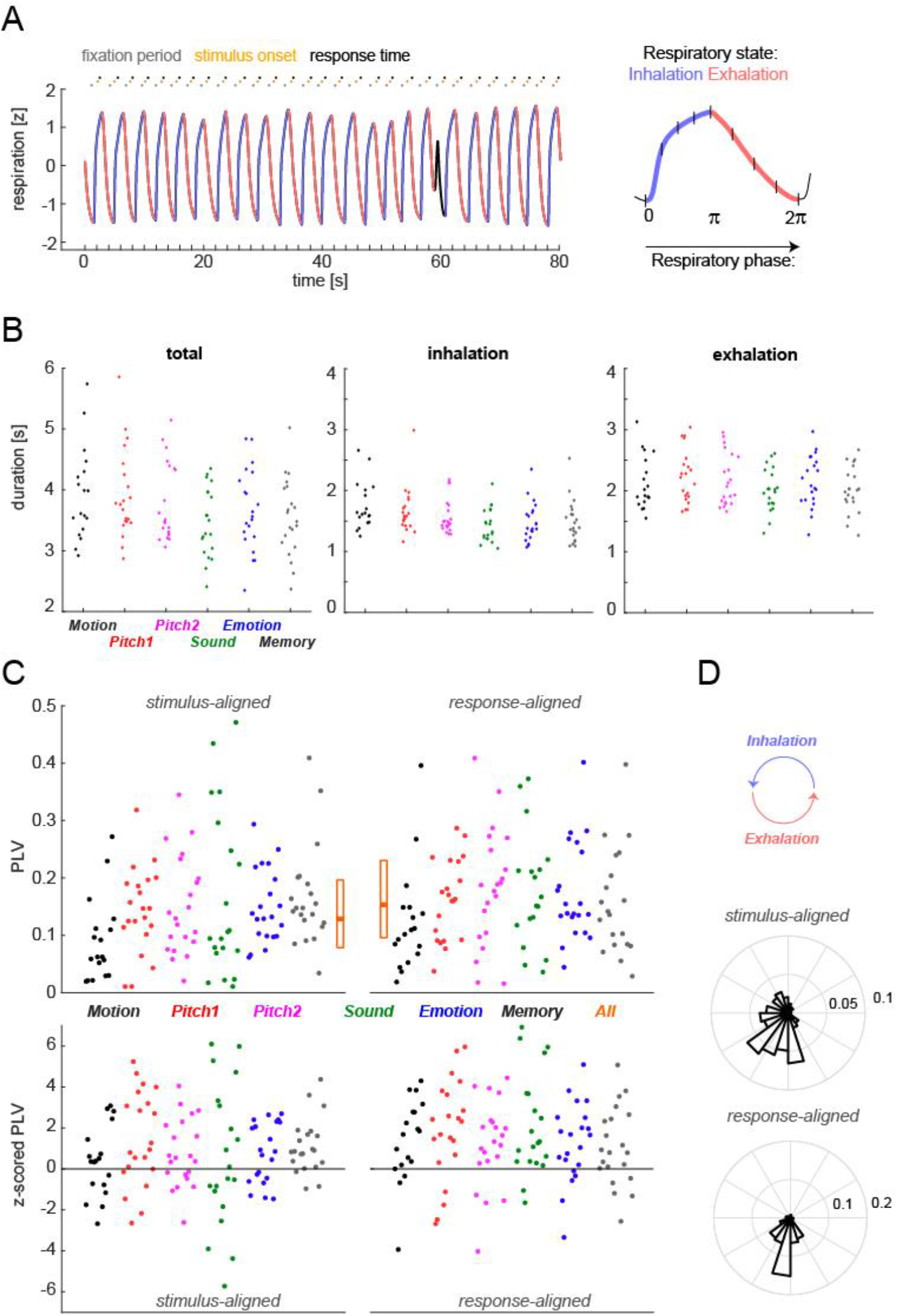
Detection of respiratory cycles and their alignment to paradigm events. **A)** Respiratory cycles were automatically segmented into inhalation(blue) and exhalation (red) periods, while atypical cycles were excluded (black). The trace shows one example respiratory signal (z-scored) together with the timing of trials for one participant in the Sound paradigm. The right panel shows how respiratory cycles were further coded as either a continuous phase variable (to test for an alignment of respiration to the paradigm); or using 4 bins phase-bins during inhalation and exhalation respectively (to test for a relation between respiration and performance). **B)** Duration of the respiratory cycles, and the inspiration and expiration states separately per paradigm (color-coded) and participant (dots indicate the participant-wise median values). **C)** Alignment of respiratory cycles to paradigm events (stimulus onset, response times). The upper panels show participant-wise phase-locking values (PLV), the lower panel the same data but z-scored to participant-specific randomization distributions (based on 2000 randomizations). Here positive values indicate stronger phase-locking than expected by chance. **D)** Distribution of participant-wise average phase values at stimulus onset or response times (derived as circular mean across trials).

To exclude atypical respiratory cycles we used two criteria. First, we compared the overall time courses of individual respiratory cycles using their mean-squared distances and excluded cycles with a distance larger than 3 standard deviations from the centroid of the participant-specific distribution. We also clustered the durations of individual inspiration and expiration epochs and excluded cycles falling outside 3 standard deviations of this centroid for each participant. These atypical cycles were excluded as they do not reflect the prototypical respiration under investigation here. Across all 122 datasets, we detected a total of 42’539 respiratory cycles, of which 41’863 were retained for analysis (98.4%). Together the excluded or undefined time points (assigned to neither inspiration or expiration) amounted to a median of 8.3% of time points (mean±s.e.m.: 9.0±1% across datasets, n=122). To link respiratory signals to behavior, we relied on the division of respiratory cycles into the two ‘states’ of inspiration and expiration. For each event of interest (stimulus onset, participant’s response times), we assigned the state as the one which prevailed in a 100ms window around the event. In addition, we subdivided each state into a continuous ‘phase’ variable (Fig. 1A, right panel). The phase was defined as linearly increasing from the beginning to the end of each state. To analyze the relation between respiration and performance this phase was binned into four equally long phase-bins (results in Figs 2-5)). To probe the alignment of the respiratory cycles to paradigm events (Fig. 1C,D), we coded the phase during inhalation (exhalation) periods as progressing from 0 to pi (pi to 2*pi), so that the full respiratory cycle could be described by a cyclic variable.

**Figure 2.**
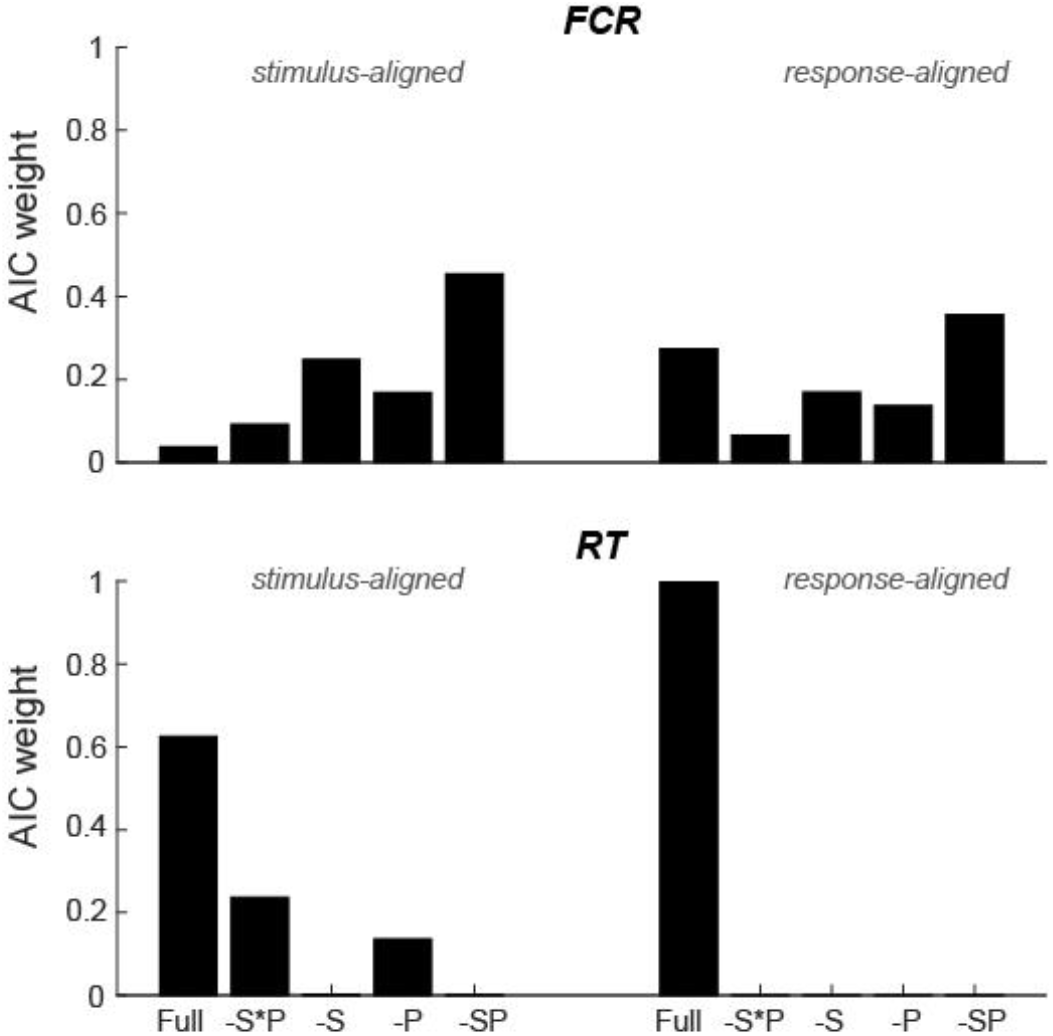
Group-level test for a statistical relation of respiratory cycles and behavior across all paradigms. Bars indicate the AIC-weights for mixed linear models prediction the dependency of the fraction of correct responses (FCR) or response times (RT) based on a model including all respiratory predictors (full), a model excluding the interaction of respiratory state and respiratory phase (-S*P; c.f. Fig. 1A for the definition of state and phase), models excluding the interaction and state (-S) or phase (-P) and a model excluding all respiratory predictions (-SP). Models were fit separately based on the respiratory variables at stimulus onset or at response times and were fit across all six paradigms and n=122 datasets (see Table 1 for model parameters).

### Statistical analysis of the alignment respiration and paradigm

To probe whether participants aligned their respiratory behavior to the experimental paradigms we proceeded in two ways. First, we coded individual respiratory cycles using their cyclic phase variable and quantified the consistency of this phase across trials using the mean-resulting vector length (phase-locking value PLV; Lowet et al. 2016). This was done for each participant and event of interest separately (stimulus onset, response time; Fig. 1C, upper panels). For comparison, we derived a surrogate distribution of phase-locking values under the null hypothesis of no alignment between respiratory trace and paradigm for each participant. This was obtained by randomly time-shifting the respiratory trace and recalculating the phase consistency 2000 times. We then z-scored, for each participant and event, the actual PLV against the randomization distribution (Fig. 1C lower panels). To test the hypothesis that participants aligned their respiratory cycle to the paradigm more than expected based on the surrogate distribution, we contrasted these z-scored PLV values against zero using sign-tests (one-tailed, using the approximate method; Rasch 1986). When testing individual paradigms, we further corrected across the 12 tests (6 paradigms, 2 alignments) using the Bejnamini & Hochberg FDR procedure (Benjamini & Hochberg 1995). We also compared the PLV between stimulus and response-aligned data using a Wilcoxon sign-rank test to probe whether the alignment strength differed between these events. In a second analysis, we probed the hypothesis that across participants the trial-averaged respiratory phase values were not distributed uniformly across the cycle (Fig. 1D). This was tested using Rayleigh’s test for non-uniformity, using the circular toolbox in Matlab (Berens 2009). To further understand whether participants specifically tended to inhale (or exhale) at the events of interest, we calculated the fraction of trials at which the respiratory state was inhalation for each event. This estimate is generally biased towards exhalation, as the overall duration of exhalation periods was longer. We again z-scored these estimates against a surrogate distribution, obtaining z-values that indicate whether individual participants tended to inhale more frequency than expected by chance.

**Table 1.**
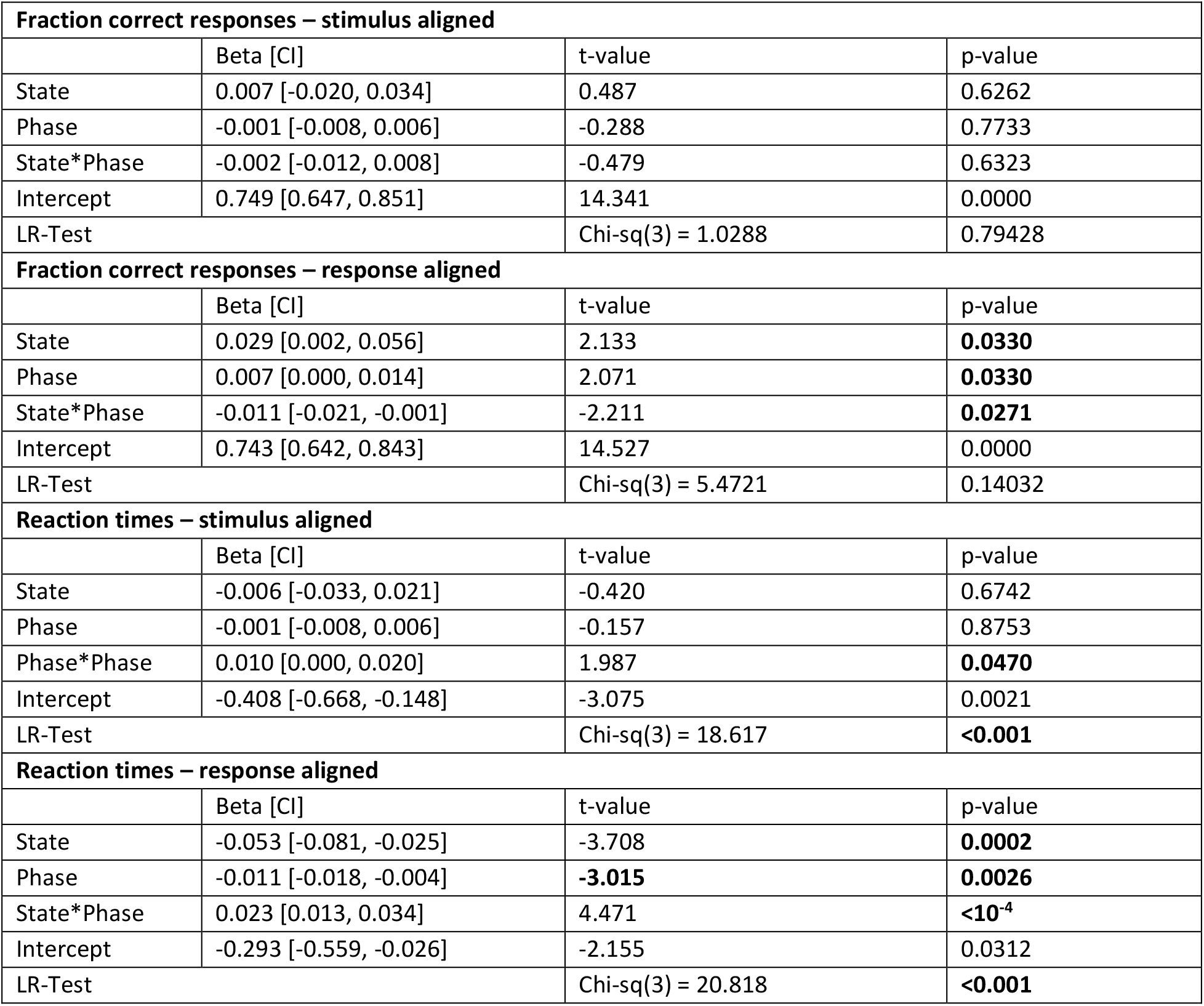
Relevance of respiratory predictors across paradigms. The relation between respiration and behavioral performance (Fraction correct responses, log-transformed reaction times) was probed using mixed linear models, which were fit separately using the state and phase of respiration obtained at stimulus onset or the response time (c.f. Fig. 1A). The table provides the predictor coefficients (incl. 95% confidence intervals), the respective t-and p-values and the result of a likelihood-ratio test comparing a model with and a model without the respiratory predictors.

### Statistical analysis relating respiratory and behavioral variables

To relate respiration and behavioral data, we first removed trials based on either atypical behavioral or respiratory data. Outliers were defined based on reaction times (RTs) being shorter than 200ms or longer than 3 standard deviations above the mean (based on log-transformed RTs). This led to the exclusion of (median) 1% of trials across datasets (mean±s.e.m: 3.5±0.8 trials; n=122). In addition, we removed trials for which the state of respiration was not defined at the events of interest. Collectively these criteria led to the exclusion of a (median) 6.5% and 7.1% of trials for stimulus-and response-aligned data (mean±s.e.m: 9.7±0.9% and 10.1±0.9% of trials, n=122). Subsequent analyses linking respiratory data to behavior focused on log-transformed reaction times (RTs; see below for rationale) and the fraction of correct responses (FCR) per condition of interest and were repeated one using the respiratory variables at stimulus onset and once at the time of participant’s response.

To code the respiratory cycle, we relied on two variables: the state of respiration (inhalation, exhalation) and the phase-bin within each state (4-levels). This coding was used to be able to differentiate inhalation from exhalation in the statistical testing, which would not be possible when focusing on a single continuous respiratory phase value. In addition, this coding normalized the overall prominence of inhalation and exhalation periods, which is important given that exhalation periods are generally longer (Fig. 1B). For each paradigm and participant, we grouped trials per stimulus level, respiratory state and phase. For each of these cells (level x state x phase) we derived the participant-averaged RT and FCR. The factor stimulus level was either the parametrically manipulated saliency of the sensory information (Motion coherence, Pitch differences, Sound signal to noise ratio), the emotion category or object novelty (Memory task). The resulting behavioral data are shown in Figure 3, averaged across phase-bins (top) or across stimulus levels (bottom). We used mixed linear models to probe whether the behavioral data indeed varied with these factors (level, respiratory state, respiratory phase, and state x phase interaction). Models were fit with participants as random effects, respiratory state as categorical effect and level and phase as linear predictors (fitlme in Matlab; using the ‘quasinewton’ method). Separate models were fit for stimulus or response aligned data. In one approach we fit models separately for each paradigm, in another approach we fit one model across all six paradigms (treating level as paradigm-specific effects). These two analyses probe whether the influence of respiratory variables i) is consistent across participants within a paradigm, or ii) is consistent across participants and also paradigms. As model outcomes we report the predictor coefficients and their 95% confidence intervals, t-and p-values (Tables 1, 2, Figure 4).

**Table 2.**
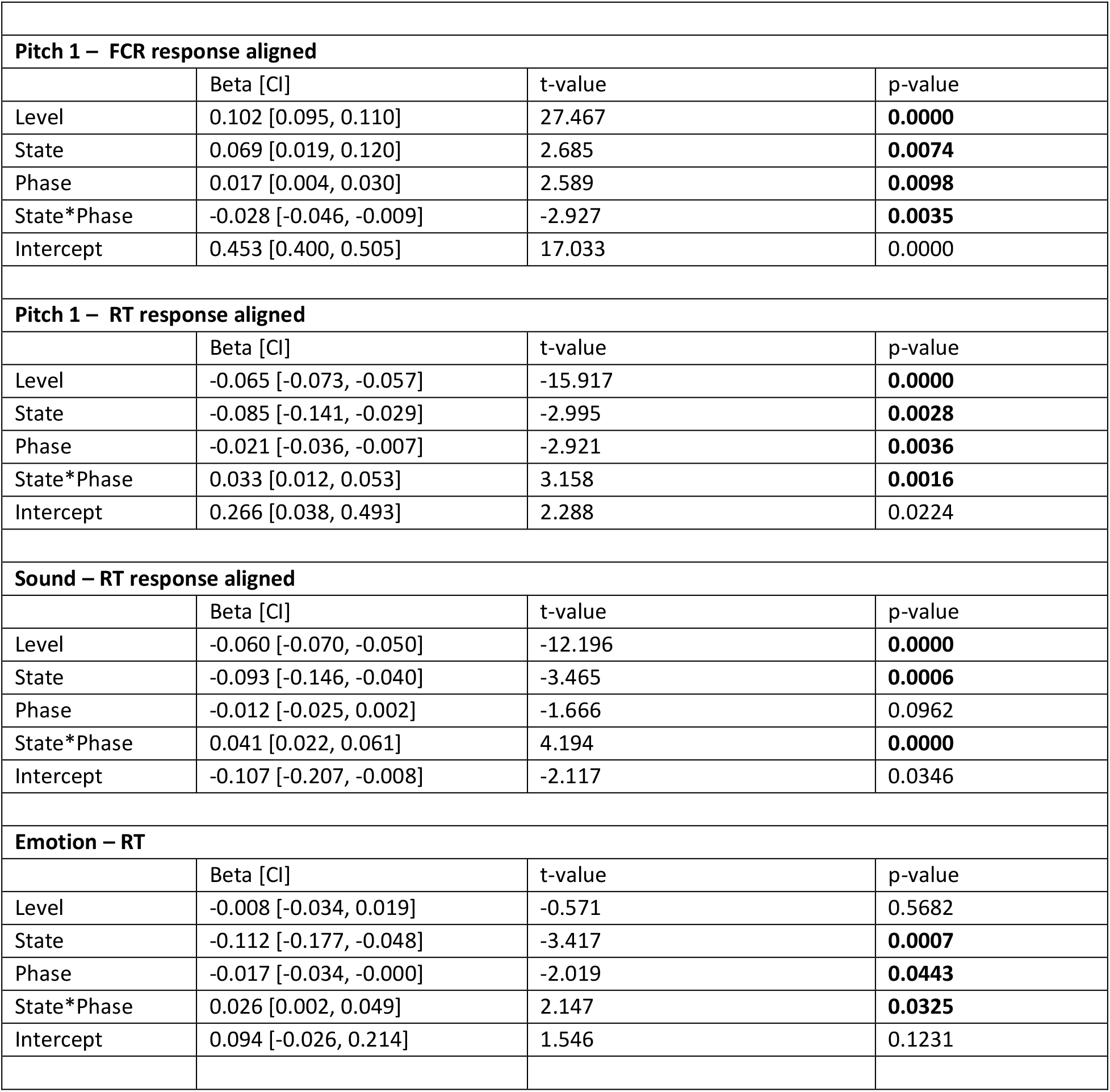
Relevance of respiratory predictors for individual paradigms. The table lists liner model results for those paradigms and alignments that returned a significant effect of respiratory variables. The results (t-statistics) for all paradigms, alignments and variables are shown in Fig. 4.

**Figure 3.**
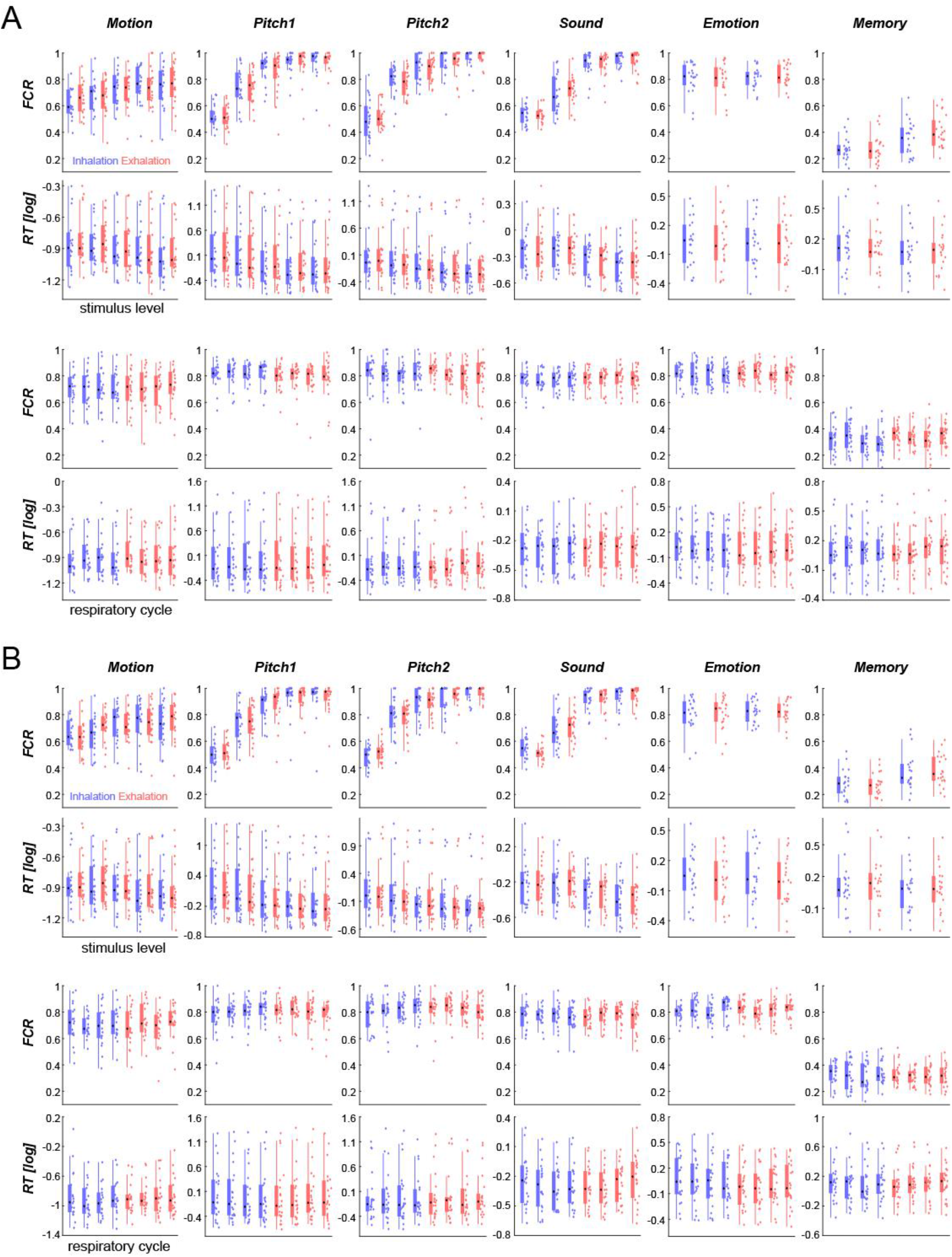
Behavioral performance for individual paradigms and participants. For each paradigm the panels show the fraction of correct response (FCR) or log-transformed reaction times (RT). The upper two panels show these as a function of respiratory state (inhalation, exhalation; color-coded) and level, the lower two panels as a function of respiratory state and phase bin (averaged over levels). A) Shows the data based on the respiratory variables at stimulus onset. B) Shows the data based on the respiratory variables at response times. Boxplots indicate the central quartiles and median (black circle), individual colored-dots the participant-wise data.

**Figure 4.**
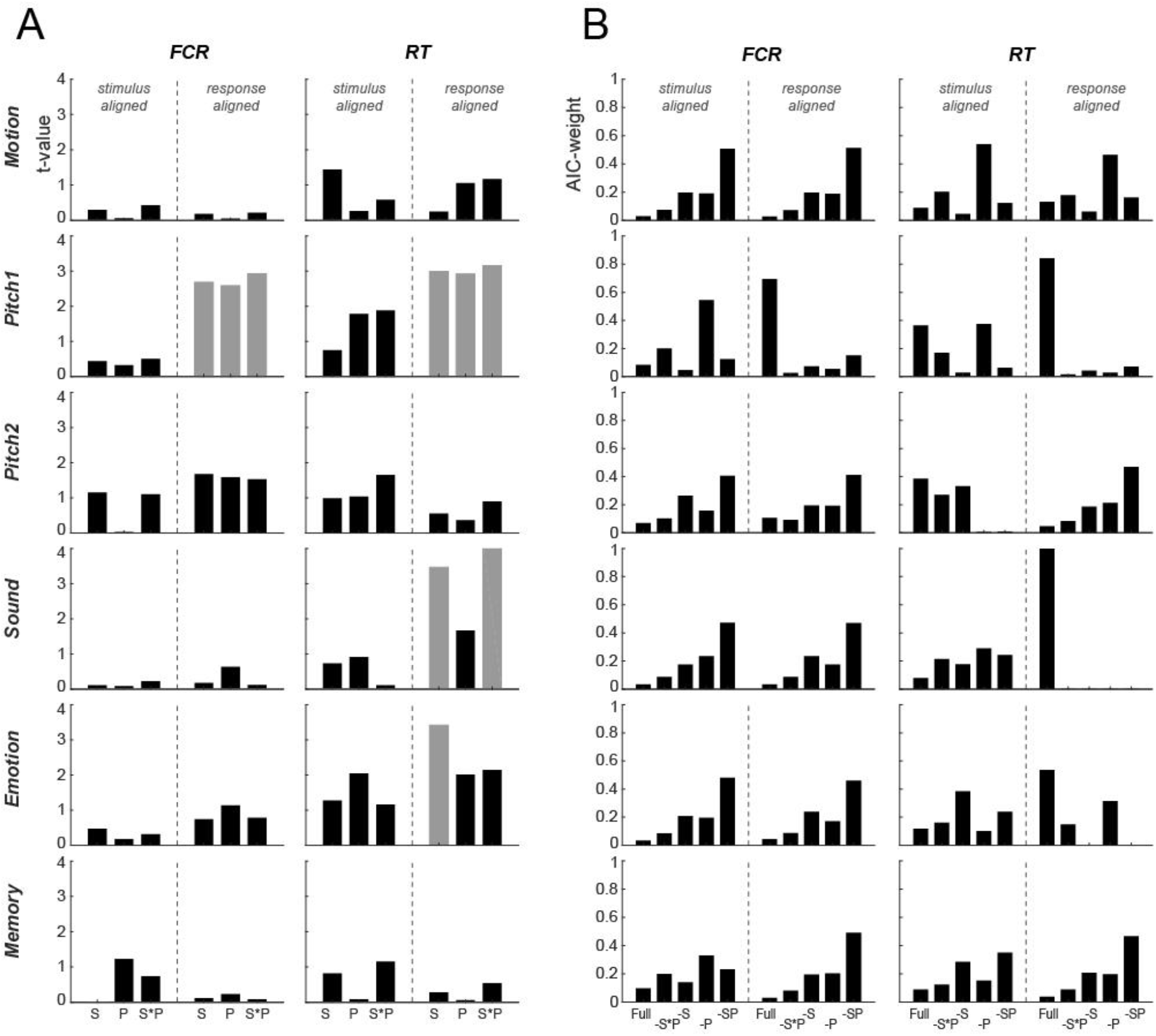
Paradigm-wise mixed-linear modelling of behavior against respiratory variables. **A)** Statistical significance (t-value) of model coefficients for linear models fit for individual paradigms and modelling the fraction of correct responses (FCR) or response times (RT) based on the respiratory variables at stimulus onset or at response times. Significant predictors are indicated in gray (p<0.01; two-sided t-test, uncorrected). Model predictors were respiratory state (S; inhalation or exhalation), respiratory phase within each state (P) and their interaction (S*P). **B)** Model comparison based on AIC-weights. Here the full model including all respiratory predictors (full) was compared to a model excluding the interaction of respiratory state and respiratory phase (-S*P), models excluding the interaction and state (-S) or phase (-P) and a model excluding all respiratory predictions (-SP).

To further substantiate whether the addition of the respiratory predictors improved the model fit, we contrasted models with and without the respiratory predictors using likelihood ratio tests (Table 1). Furthermore, we systematically contrasted the explanatory power of models including or excluding individual respiratory predictors based on their respective log-likelihoods: we fit models including all three respiratory predictors, a model excluding their interaction, two models excluding the interaction and either state or phase, and a model excluding all three respiratory predictors. We used the Akaike information criterion to derive the conditional probability of each model given the data using the Akaike weights (Wagenmakers & Farrell 2004).

We based the analysis of reaction times on the log-transformed data as a preliminary analysis had revealed that this allowed the best description of the expected effects of the stimulus manipulations (levels). Specifically, for the four tasks with expected effects of stimulus levels (Motion, Pitch 1&2, Sound) we applied the above described linear mixed models to only the factor level, after deriving the level-specific average reaction time using three different coding schemes (Lo & Andrews 2015): the raw data, the log-transformed data, or the inverse-transform. This revealed that the log-transformed data allowed the best linear-model-based description based on the summed AIC across all four paradigms (AICs for raw values, log-and inverse-transform: -306, -526, -351), although all three coding schemes allowed capturing the effect of level well (adjusted R^2^: 0.96, 0.96, 0.95).

### Analysis of effect sizes

To quantify the effect size of the dependency of behavioral data on respiratory variables (Fig. 5), we first described the group-level median data using a rhythmic dependence on respiratory time, after averaging out the factor level. For this we compared the explanatory power (log-evidence) of models involving distinct timescales between 0.5 and 3 Hz per respiratory cycle, and derived the best-fitting time-scale (Fig. 5; gray curves). We then used these rhythmic descriptors to quantify an effect-size of how much reaction times or the FCR vary along the respiratory cycle: for each paradigm and factor of interest we computed the difference in reaction times (or FCR) between the bins with positive or negative rhythmic components. Note that this fitting of rhythmic models was only used to decide on which phase-bins to contrast, not to substantiate the statistical significance of a rhythmic modulation. These effect-sizes were derived for each participant individually and reflect the expected modulation of RT (of FCR) under the assumption that the time course of such modulation is the same across all participants. Group-level Effect sizes were quantified using their mean, median and Hedge’g. The 95% confidence intervals for the mean were obtained from the respective t-distribution, for the median they were obtained using the Harrell-Davis estimator based on the ‘matrogme’ Matlab package (Wilcox 2012; Rousselet et al. 2017), and for Hedges’g they were obtained using bootstrapping using the Effect size toolbox in Matlab based on 2000 bootstrap samples (Stüttgen, 2011).

**Figure 5.**
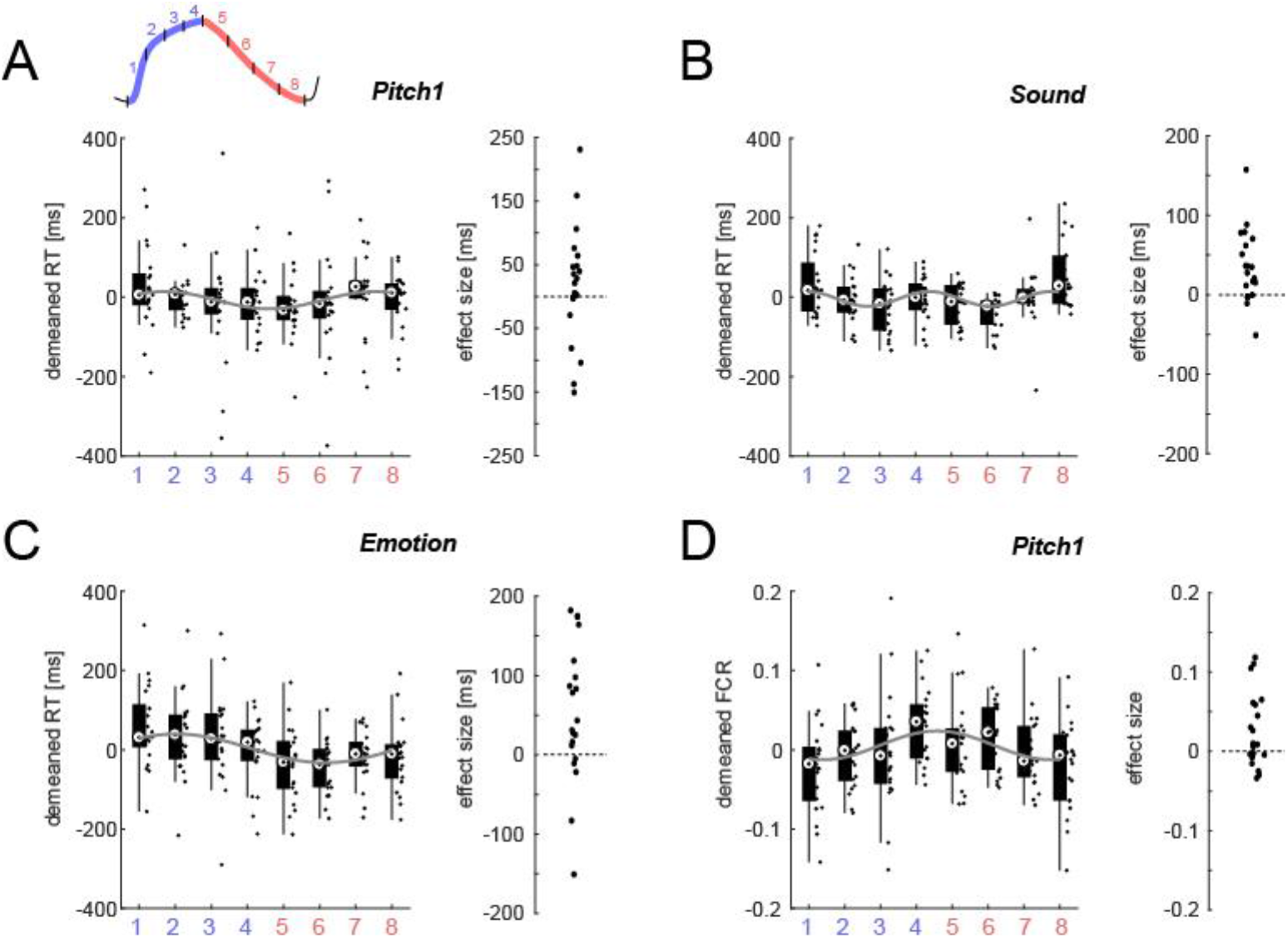
Effect sizes of behavioral changes along the respiratory cycle. For those paradigms with significant effects in linear models (c.f. Table 2) this figure shows the respective variation in reaction times (**A-C**) or the fraction of correct responses (**D**) along the respiratory cycle (inset in Panel A). For display purpose, the individual participant data were mean-subtracted. The group-level median data were fit using a rhythmic model (gray line), based on which we determined those bins with positive and negative RT (or FC). We then derived a measure of effect size as the participant-wise difference in the data between positive and negative bins. This procedure ensures that effect sizes are derived under the assumption of a fixed respiratory pattern across participants within a paradigm. The right-hand panels indicate the participant-wise (dots) effect sizes.

## Results

### Properties of respiratory cycles

We collected behavioral data during six paradigms involving the detection or discrimination of either visual or acoustic stimuli or testing participant’s memory of these. This resulted in a total of 122 datasets. During each of these experiments participant’s respiration was recorded using a thermistor inserted into a face mask. This provided high resolution traces reflecting the temperature changes induced by inhalation and exhalation (Fig. 1A). These were automatically segmented into individual respiratory cycles. Across datasets we detected a total of 42’539 cycles, of which 98.4% were retained for analysis. The time course of individual cycles was split into two variables for analysis: the respiratory state was defined as either inhalation or exhalation (Fig. 1A; color-coded); the time within each state was divided into a phase variable, which increased linearly from the start to the end of each state (Fig. 1A; right panel).

Across participants the median duration of respiratory cycles was 3.57 s ([25th, 75th percentiles]: [3.25, 4.15] s, n = 122; durations were averaged within participants across detected cycles, Fig. 1B), corresponding to a respiratory frequency of ∼0.3 Hz. As expected, inhalation states were shorter than exhalation states (inhalation: 1.49 s [1.34, 1.69] s; exhalation: 2.03 s [1.82, 2.41] s). Figure 1B provides the respective durations per participant and paradigm.

### Respiratory cycles are aligned to paradigm events

Previous studies suggested that participants may potentially adjust their respiratory behavior to an experimental paradigm, for example by aligning their respiratory cycle with expected events such as stimulus presentation or their responses (Perl, et al., 2019). We hence probed for signatures of such behavioral strategies in the present data. In four of the paradigms tested here, the stimuli were presented at pseudo-random intervals following a fixation onset (0.4 - 1 s uniformly distributed) or following participants self-initiated trial-start (Pitch 2, 0.3 - 0.6 s, uniform delay). Alternatively, participants may have aligned their respiratory pattern to their responses.

We tested for such an alignment by computing for each participant the phase-locking value (PLV) characterizing the consistency of the respiratory cycle across trials (Fig. 1C, upper panels). We then compared the resulting PLV within participants to a surrogate values obtained under the null hypothesis of no alignment between respiratory trace and the experimental paradigm (Fig. 1C, lower panels). We first tested the hypothesis that the alignment in the actual data was stronger than in the surrogate data: across paradigms this was the case both for stimulus onset and at participant’s response times (n=122; one-sided sign-tests, stimulus onset Z_sign-test_=3.5, p=0.004, response time Z_sign-test_=6.8, p<10^−4^). Repeating this test for individual paradigms returned a significant effect for each paradigm in the response-aligned data (corrected using the Bejnamini & Hochberg FDR procedure across the 12 tests, minimal Z_sign-test_=2.1, at least p_corr_ <0.02) and for the memory paradigm also in the stimulus-aligned data (Z_sign-test_=3.0, p_corr_ = 0.004).

Inspecting the individual participant data revealed that for many participants the observed PLV values were significantly stronger than expected based on their individual surrogate data (individual z-scored PLVs larger than z=3: 19 of 122 for stimulus onset and 28 of 122 for response time; Fig. 1C, lower panel). This difference between stimulus and response-aligned data seems to suggest that the alignment was stronger when analyzed contingent on response time. Indeed, a comparison of the actual PLVs between stimulus and response aligned data revealed a significant difference (Wilcoxon sign-rank test, Z_sign-rank_=-4.2, p<10^−4^): the PLV values were higher at response time (median and [25th, 75th percentiles]; 0.13 [0.08, 0.2] and 0.15 [0.1, 0.23] for stimulus onset and response time; Fig. 1C, upper panel).

To understand how precisely respiratory cycles were aligned to the paradigm, we investigated the trial-averaged respiratory phase angles (coded as 0 to pi for the inhalation period and pi to 2 pi for exhalation Fig. 1D). The average phase angles were highly clustered across participants, and significantly deviated from a null hypothesis of a uniform distribution (Rayleigh tests, Z_Rayleigh_ =25 and Z_Rayleigh_ =72, both p<10^−4^ for stimulus and response-aligned data). These data visualize the stronger alignment of respiratory behavior around response time across participants and support the conclusion that participants systematically aligned their respiratory behavior to the paradigm. However, because exhalation periods were generally longer, the average phase angle in Fig. 1D may confound the prominence of exhalation periods with an alignment specifically emphasizing exhalation around response times. In a final analysis we hence calculated the fraction of trials featuring an inhalation state at the events of interest, and z-scored this within participants against the individual expected proportion based on surrogate data. We then asked whether the median proportion of excess (vs. surrogate) inhalation states deviated from zero. Across datasets, this revealed a significant shift towards inhalation near stimulus onset (median z=0.64, Wilcoxon sign-rank test, Z_sign-rank_ =3.2, p=0.001) and a shift towards exhalation near response times (median z=-0.47, Z_sign-rank_=-0.32, p=0.001).

### Behavioral data covary with the respiratory cycle

We then asked whether participants behavioral performance revealed evidence for a systematic variation along the respiratory cycle. For each paradigm, we focused on the fraction of correct responses (FCR) and participant’s reaction times (RTs) and related these to the respiratory variables state and phase. We tested for such relations using linear mixed models, separately for the respiratory variables obtained at stimulus presentation or response time. In a first analysis, we probed for such a relation across all six paradigms, hence testing the hypothesis that behavioral performance varies with respiration consistently across experiments. Across datasets we included (based on acceptable behavioral and respiration data) around 93% of all trials in the analysis (c.f. Material and Methods). We first grouped trials based on the respiratory variables (state, phase, their interaction) and based on stimulus level (depending on paradigm). We then computed for each participant the FCR across trials per group and the trial-average log-transformed RTs. The model was fit across participants (random effects) to predict FCR or RT based on stimulus level (fixed effect per paradigm) and respiration status (fixed effects).

Across the four analyses (FCR and RT; stimulus and response alignment) we found clear evidence that RTs were related to respiration. For RTs, the interaction of respiratory state and phase was a significant predictor in the stimulus-aligned data (p<0.05; see Table 1 for detailed results) and all factors were highly significant for the response-aligned data (p < 0.001; see Table 1). To substantiate this observation, we contrasted the GLMMs with and without the respiratory predictors. Likelihood-ratio tests returned a significant improvement in model fits for both alignments when including respiration (p<0.001; see Table 1). In addition, we also compared models featuring only individual respiratory variables and a reduced model excluding all respiratory predictors. For each model we derived its conditional probability within the range of tested models based on Akaike weights (Fig. 2). This yielded clear evidence for an association of respiratory variables and RTs in the response aligned data (AIC-weight for the full model was 0.998) but not the stimulus-aligned data (AIC-weight of full model 0.62). In sum, these results show that participant’s reaction times are associated with the respiratory cycle near the time of these responses.

For the FCR, the respiratory predictors were marginally significant in the response-aligned (p<0.05; see Table 1) but not the stimulus-aligned data. Likelihood-ratio tests returned no evidence for an improvement in model fit when including respiratory predictors (p>0.05; Table 1) and the comparison of individual models also returned no evidence for an association of FCR and respiration based on AIC-weights (Fig. 2).

### Paradigm specific results

This results point to a systematic relation between respiration and reaction times but not response accuracy. Yet, probing for generic and paradigm-independent effects may also obscure more specific results, such as for example stronger effects for the paradigms involving medial temporal brain structures (Emotion, Memory), for which previous studies makes clear predictions.

We hence repeated the analyses for individual paradigms. Figure 3 displays the individual and group-level behavioral data showing the effects of stimulus level and respiratory state (upper two panels in A, B) and showing the effects of the respiratory cycle averaged across levels (lower two panels in A, B). For each paradigm, we again used linear mixed models to probe the relation between behavior and respiration. As expected, for each paradigm manipulating stimulus saliency in a parametric fashion (Motion, Pitch1, Pitch 2, Sound), the effect of stimulus level was significant for both FCR and RTs and for both alignments (all t-values > 4.5, p < 10^−4^). For the discrimination of emotional faces (Emotion), the effect of emotion was not significant for neither FCR nor RTs, and for neither alignment (all t< 1.2, p> 0.25). For the memory test, the effect of novelty was significant for FCR (both alignments t> 6.1, p < 10^−4^) and for RTs (t= 3.9 and 2.5, p = 0.0001 and 0.012 for stimulus and response alignment).

The effects of the respiratory variables are summarized in Figure 4A, which shows the respective t-statistics and significances for individual predictors (color-coded). Figure 4B shows the conditional probabilities (AIC-weights) for the full model and models excluding individual respiratory predictors. When modelling FCR, respiration was a significant predictor in the response aligned data for Pitch 1 (see Table 2 for details); in this case the full model including all respiratory predictors had the highest probability in this case (AIC-weight 0.69). For all other paradigms and alignments, the model excluding all respiratory predictors generally had the highest conditional probability when modelling FCR (AIC-weight; Fig. 4B).

When modelling RTs, the respiratory variables at the response time were significant predictors for three paradigms: pitch discrimination (Pitch1; c.f. Table 2 for details), sound detection (Sound) and the discrimination of emotions (Emotion). For these paradigms the full model including all respiratory predictors also had the highest conditional probabilities (AIC-weights 0.84, 0.99 and 0.53; Fig. 4B). In contrast, the respiratory variables derived at stimulus onset were not significant predictors for RTs in any paradigm (Fig. 4A). Collectively these results confirm a significant relation between respiration at the time of participant’s responses and behavior performance across a number of paradigms.

One experiment involved the detection (rather than discrimination) of stimuli and was additionally analyzed using the framework of signal detection theory. For the Sound paradigm, we calculated hit-and false alarm-rates as well as d’ and bias (n=22). The linear models for none of these parameters revealed a significant effect for any of the respiratory features (t<1.1, p>0.26) and the AIC-weights did not show evidence for models including respiration to perform better than those without (no model exceeded an AIC-weight of 0.5). Hence, we did not find evidence for any change in response behavior with respiration in this paradigm other than of the reaction times.

### How much does behavior co-vary with the respiratory cycle

We investigated the relation between the respiratory cycle and behavior for those paradigms with significant effects in more detail. Figure 5 visualizes individual participant’s reaction times (in milliseconds) and the fraction of correct responses (for Pitch 1) along the respiratory cycle. To obtain effect sizes we subtracted RTs (FCR) of those four bins with higher performance from those four with lower performance. To determine those bins under the assumption of a consistent group-level effect, we first derived the best-fitting rhythmic description of the group-level data using a sinusoidal dependence on respiratory time (Fig. 5, gray lines). Based on this rhythmic description we decided on how to group bins to determine effect sizes (see legend Fig. 5). For comparison with previous work we quantified the effect sizes and their confidence intervals using three descriptors. The group-level mean effect sizes for RTs were 17 ms, 36 ms and 67ms (95% confidence intervals [-23, 58]ms, [15, 57]ms, and [18, 114]ms respectively; n = 21, 20, 20). The median effect sizes for RTs were 20 ms, 30 ms, 54 ms (with 95% bootstrap confidence intervals of [-8, 46]ms, [12, 55]ms, and [17, 109]ms). When quantified using Hedges’ g the effects were 0.19, 0.79 and 0.62 (95% bootstrap confidence intervals [-0.22, 0.67], [0.42, 1.3], [0.22, 1.14]). The FCR for Pitch 1 varied by a mean of 0.025 ([0.0045, 0.046], n=21), a median of 0.013 ([-0.002, 0.04]), with Hedge’s g of 0.55 ([0.19, 0.95]).

## Discussion

We asked whether and by how much participant’s performance in diverse perceptual tasks varies along the respiratory cycle. Across six sensory detection, discrimination and a memory task we found that participants tended to align their respiratory cycle to the experimental paradigm. Their reaction times consistently and significantly varied along the respiratory cycle, while response accuracies varied little. This covariation of human behaviour with the respiratory cycle was stronger when analysed contingent on the state of respiration around participant’s responses than around stimulus onset. These results support an intricate, and possibly causal, relation between respiration and sensory-cognitive function and show that the respiratory state may be an important factor to consider when investigating cognition and its neural underpinnings.

### Effect sizes of respiratory modulation

A number of studies have shown that human performance in sensory (Flexman et al. 1974; Gallego et al. 1991), mental (Arch & Craske 2006; Homma & Masaoka 2008; Vlemincx et al. 2010; Zelano et al. 2016; Perl et al. 2019), memory (Arshamian et al. 2018; Huijbers et al. 2014; Nakamura et al. 2018; Zelano et al. 2016) or motor tasks (Park et al. 2020; Li & Rymer 2011) varies along the respiratory cycle. Yet, obtaining a comprehensive picture has remained difficult for a number of reasons. Different studies used different technical and statistical approaches to detect variations in behavioral performance along the respiratory cycle, making it difficult to compare effects across experimental tasks. Furthermore, many studies instructed participants to a specific type of respiration, thus possibly biasing attention to their own respiration (Varga & Heck 2017; Herrero et al. 2018; Vlemincx et al. 2010). In contrast, we systematically assessed the relation between task performance and respiration across six tasks and in a large participant sample (n=122), that were performing these laboratory tasks without receiving specific instructions as how to breath.

The covariation of respiratory cycle and behavior was more prevalent when analyzed based on the respiratory cycle extracted around participant’s responses, both when combining the data across paradigms and when considering them individually. Possibly, this relates to a tendency of participants to align their respiration around their responses rather than to the more uncertain stimulus onset. This view is consistent with a causal and mechanistic relation between respiration and reaction times, as discussed below, although determining the causality and its direct neural mechanisms requires further work.

While previous studies often focused on detecting statistically significant relations between behavior and respiration, we investigated the respective effect sizes and their associated uncertainty. This is particularly important to understand by how much task performance changes along the respiratory cycle. In our data changes in response accuracy were either statistically insignificant (5 out of 6 paradigms) or were small (1.3% of correct responses). Importantly, these effects were associated with confidence intervals including zero and hence do not allow a firm conclusion on a positive effect. This observation is consistent with previous work reporting no significant effect on discrimination performance in the same emotion task as used here (Zelano et al. 2016), were our data replicate the absence of a significant effect reported previously. Contrasting this, one study reported significant differences in a visuo-spatial shape discrimination task on the order of 5% and replicated in this in two groups of participants. However, the respective confidence intervals reported in that study included zero casting doubts on the existence of a clear effect (Perl et al. 2019, experiment 4). Furthermore, the same study found no significant relation to task performance in a lexical decision task the (Perl et al. 2019). A recent study looking at a visual discrimination task quantified the change in psychometric threshold as a function respiratory cycle and reported shifts on the order of 5% of the respective thresholds, but no confidence for these effects was reported (Kluger and Gross 2020). A notable deviation from this picture comes in an older study investigating the detection of visual signals, which reported a 22% difference in detection rates between inhalation and exhalation periods (95%-Students CI derived from the data reported in that paper: [19-26]%; (Flexman et al. 1974). In sum, we take this body of work to suggest that any relation between response accuracy and respiration is small and most likely not consistent across sensory-cognitive domains.

In contrast, reaction times covaried significantly with the respiratory cycle in 3 out of 6 paradigms tested here. Based on the associated confidence intervals, the effects sizes for RTs fall in between 10 ms to 110ms, with median values between 20ms and 50ms. This order of effect is consistent with previous studies reporting a change in RTs for a sound detection task with a mean effect size of 17ms for spontaneous breathing and 61ms for controlled breathing (Gallego et al. 1991; confidence interval could not be extracted as no data variability was reported). Similarly, a study on the discrimination of emotional faces reported an effect size of 16ms (95% CI estimated from the reported data: [4-29]ms; Zelano et al. 2016 Emotion task with nasal breathing). In some studies, even stronger effects were found for controlled or deep respiration, as well as forced oral respiration, suggesting that attention to respiration can further enhance these effects (Gallego et al. 1991; Zelano et al. 2016; Li & Laskin 2006). Together these studies suggest that reaction times on sensory-cognitive tasks indeed vary systematically with the state of respiration.

To put the effect size of changes in reaction times along the respiratory cycle in context, we here compare these to other and well-established changes in reaction times in sensory-cognitive paradigms. For example, a meta-study on the effect of transcranial magnetic stimulation in attention and memory tasks (Beynel et al. 2019) reported an overall effect size of 0.24 (Hedge’s g; 95% confidence interval [0.05, 0.42]). Similarly, a meta-analysis of the effects of transcranial direct current stimulation on inhibitory control reported even smaller effect sizes, with an average of 0.10 (Hedge’s g; 95% confidence interval [0.02, 0.21] and a largest effect size per individual study of up to 0.83 (Schroeder et al. 2020). Compared to such effects of brain stimulation, the effect sizes of respiratory-related changes in reaction times are very comparable (Hedge’s g between 0.19 and 0.79). Comparable or sizes were reported for cognitive interference effects in clinical populations (average of Cohen’s d of 0.67 in (Szucs & Ioannidis 2017) and in a larger meta-analysis of a large body of cognitive neuroscience and psychology literate (average d of 0.66; Szucs & Ioannidis 2017). Hence, changes in reaction times along the respiratory cycle may be well comparable to other changes in reaction times induced by typical sensory-cognitive paradigms.

### Respiration as potentially confounding factor in sensory-cognitive studies

These results also suggest that respiration may act as a confounding factor in sensory-cognitive paradigms, in particular as participants may align their respiratory pattern to specific experimental manipulations. Neuroimaging studies have considered respiration as a major confounding factor for a long time, via the associated movements of body and brain (Myllylä et al. 2017; Fair et al. 2020). However, respiration may also affect heartrate (Bernston et al. 1993) and thereby neuroimaging signals indirectly: the visibility of ECG-related signals in electro-magnetic brain measurements and pulsation-related effects can distort these in a respiration-specific manner. The sensation of one owns heartbeat is reflected in neural activity in the brain (Petzschner et al. 2019; Montoya et al. 1993), hence leaving a route for respiration to affect brain activity also via other interoceptive mechanisms. Collectively, this suggests that respiration may not only be a potential confounder for measurements of brain activity but may also causally shape, directly or indirectly, brain activity and behaviour (Heck et al. 2017; Varga & Heck 2017). One could imagine that such respiration-mediated effects go hand in hand with other effects by which the brain predicts upcoming events in an experimental paradigm, or their perceptual salience, as these can provide reliable cues for respiration to align with.

### Causal roles of respiration

Measurements of brain activity have shown that brain activity aligns to the respiratory cycle not only within olfactory structures, but also in brain regions connected with the olfactory system such as the limbic system (Boiten et al. 1994), and even in more distant regions such as parietal or prefrontal cortex (Tort et al. 2018; Jung et al. 2019; Kluger & Gross 2020; Herrero et al. 2018). While the specific impact of respiratory signals in higher association regions remains to be fully understood, recent studies have shown that such respiratory-modulations also extend into the motor system, offering one of potentially several pathways by which reaction times may be altered in a respiratory-specific manner (Park et al. 2020; Kluger and Gross 2020).

A second pathway for respiration to shape perception may be via the modulation of neural excitability in sensory regions. An MEG study showed that parieto-occipital alpha band activity modulates along the respiratory cycle (Kluger & Gross 2020). In that study the significant respiration-alpha coupling temporally preceded performance changes in perceptual accuracy that were also related to respiration. Alpha band activity reflects the excitability of cortical circuits (Mumford 1992; Haegens et al. 2011; Kayser et al. 2015) and is an omnipresent predictor of perceptual performance, reaction times or metacognition in sensory-cognitive tasks (Busch & VanRullen 2010; Henry & Obleser 2012; Iemi et al. 2017). In particular, studies have shown a correlation between alpha power more direct measures of neural activity (Varga & Heck 2017; Mumford 1992; Fontanini & Bower 2006). If respiration indeed shapes parietal alpha band activity (Hsu et al. 2020), it is not surprising hence, that respiration shapes perceptual performance across a range of tasks. Given that alpha band activity is also known to be influenced by factors such as spatial or cross-modal attention (Klimesch 2012; Thut et al. 2006), temporal expectation (Foster et al. 2017) or idiosyncratic biases (Grabot & Kayser, 2020), it remains unclear to what degree any respiratory influence becomes visible in each individual experimental paradigm. Our comprehensive analysis across six tasks suggests that effects on response speed may be rather.

## Conclusion

It is becoming clear that the sensory-cognitive performance of animals and humans can only be understood with a comprehensive view on the both the external environment and the intrinsic state. The networks receiving external evidence and those serving interoception are intricately linked (Ondobaka et al. 2017), and studies show that the homeostatic state and visceral body functions such as heartbeat bear an influence on perceptual reports. For example, the endocrine system can shape perceptual thresholds via levels of glucocorticoids (Obleser et al. 2021), while the heartbeat affects perception not only in the somatosensory domain (Al et al. 2020) but also influences value-based decisions (Azzalini et al. 2020) and can guide visual exploration search (Galvez-Pol et al. 2020). All in all, this supports the notion of active sensing, whereby the integration and exploitation of sensory information is controlled by a combination of external, intrinsic and metacognitive processes, which often operate on a seemingly rhythmic basis. (Corcoran et al. 2018).

## Acknowledgements

We would like to thank Miriam Niemeier for help with collecting the data.

## References

Al, E. et al., 2020. Heart–brain interactions shape somatosensory perception and evoked potentials. PNSA, 5, Volume 117, pp. 10575–10584.

Arch, J. J. & Craske, M. G., 2006. Mechanisms of mindfulness: Emotion regulation following a focused breathing induction. Behaviour Research and Therapy, 12, Volume 44, p. 1849–1858.

Arshamian, A., Iravani, B., Majid, A. & Lundström, J. N., 2018. Respiration Modulates Olfactory Memory Consolidation in Humans. The Journal of Neuroscience, 10, Volume 38, p. 10286–10294.

Azzalini, D., Buot, A., Palminteri, S. & Tallon-Baudry, C., 2020. Responses to heartbeats in ventromedial prefrontal cortex contribute to subjective preference-based decisions.

Benjamini, Y. & Hochberg, Y., 1995. Controlling the False Discovery Rate: A Practical and Powerful Approach to Multiple Testing. Journal of the Royal Statistical Society. Series B (Methodological), 1, Volume 57, pp. 289–300.

Berens, P., 2009. CircStat: AMATLABToolbox for Circular Statistics. Journal of Statistical Software, Volume 31.

Berntson, G. G., Cacioppo, J. T. & Quigley, K. S., 1993. Respiratory sinus arrhythmia: Autonomic origins, physiological mechanisms, and psychophysiological implications. Psychophysiology, 3, Volume 30, p. 183–196.

Beynel, L. et al., 2019. Effects of online repetitive transcranial magnetic stimulation (rTMS) on cognitive processing: A meta-analysis and recommendations for future studies. Neuroscience and Biobehavioral Reviews, 12, Volume 107, pp. 47–58.

Boiten, F. A., Frijda, N. H. & Wientjes, C. J. E., 1994. Emotions and respiratory patterns: review and critical analysis. International Journal of Psychophysiology, 7, Volume 17, p. 103–128.

Busch, N. A. & VanRullen, R., 2010. Spontaneous EEG oscillations reveal periodic sampling of visual attention. Proceedings of the National Academy of Sciences, 8, Volume 107, p. 16048–16053.

Corcoran, A. W., Pezzulo, G. & Hohwy, J., 2018. Commentary: Respiration-Entrained Brain Rhythms Are Global but Often Overlooked. Frontiers in Systems Neuroscience, 6.Volume 12.

Ebner, N. C., Riediger, M. & Lindenberger, U., 2010. FACES—A database of facial expressions in young, middle-aged, and older women and men: Development and validation. Behavior Research Methods, 2, Volume 42, p. 351–362.

Fair, D. A. et al., 2020. Correction of respiratory artifacts in MRI head motion estimates. NeuroImage, 3, Volume 208, p. 116400.

Flexman, J. E., Demaree, R. G. & Simpson, D. D., 1974. Respiratory phase and visual signal detection. Perception & Psychophysics, 3, Volume 16, p. 337–339.

Fontanini, A. & Bower, J. M., 2006. Slow-waves in the olfactory system: an olfactory perspective on cortical rhythms. Trends in Neurosciences, 8, Volume 29, p. 429–437.

Foster, J. J. et al., 2017. Alpha-Band Oscillations Enable Spatially and Temporally Resolved Tracking of Covert Spatial Attention.

Gallego, J., Perruchet, P. & Camus, J.-F., 1991. Assessing Attentional Control of Breathing by Reaction Time. Psychophysiology, 3, Volume 28, p. 217–224.

Galvez-Pol, A., McConnell, R. & Kilner, J. M., 2020. Active sampling in visual search is coupled to the cardiac cycle. Cognition, 3.Volume 196.

Grabot, L. & Kayser, C., 2020. Alpha Activity Reflects the Magnitude of an Individual Bias. The Journal of Neuroscience, April, p. 3443–3454.

Grimaud, J. & Murthy, V. N., 2018. How to monitor breathing in laboratory rodents: a review of the current methods. Journal of Neurophysiology, 8, Volume 120, p. 624–632.

Haegens, S. et al., 2011. -Oscillations in the monkey sensorimotor network influence discrimination performance by rhythmical inhibition of neuronal spiking. Proceedings of the National Academy of Sciences, 11, Volume 108, p. 19377–19382.

Heck, D. H. et al., 2016. Cortical rhythms are modulated by respiration. Cold Spring Harbor Laboratory, April.

Heck, D. H. et al., 2017. Breathing as a Fundamental Rhythm of Brain Function. Frontiers in Neural Circuits, 1.Volume 10.

Henry, M. J. & Obleser, J., 2012. Frequency modulation entrains slow neural oscillations and optimizes human listening behavior. PNAS, Volume 109, pp. 20095–20100.

Herrero, J. L. et al., 2018. Breathing above the brain stem: volitional control and attentional modulation in humans. Journal of Neurophysiology, 1, Volume 119, p. 145–159.

Homma, I. & Masaoka, Y., 2008. Breathing rhythms and emotions. Experimental Physiology, 8, Volume 93, p. 1011–1021.

Hsu, S.-M., Tseng, C.-H., Hiseh, C.-H. & Hsieh, C.-W., 2020. Slow-paced inspiration regularizes alpha phase dynamics in the human brain. JNP, January.

Huijbers, W. et al., 2014. Respiration phase-locks to fast stimulus presentations: Implications for the interpretation of posterior midline “ deactivations” . Human Brain Mapping, 4, Volume 35, p. 4932–4943.

Iemi, L., Chaumon, M., Crouzet, S. M. & Busch, N. A., 2017. Spontaneous Neural Oscillations Bias Perception by Modulating Baseline Excitability. Journal of Neuroscience, 1, Volume 37, pp. 807–819.

Jung, F. et al., 2019. Respiration competes with theta for modulating parietal cortex neurons. 7.

Kayser, C. et al., 2015. Rhythmic Auditory Cortex Activity at Multiple Timescales Shapes Stimulus-Response Gain and Background Firing. Journal of Neuroscience, 5, Volume 35, p. 7750–7762.

Kayser, S. J., McNair, S. W. & Kayser, C., 2016. Prestimulus influences on auditory perception from sensory representations and decision processes. Proceedings of the National Academy of Sciences, 4, Volume 113, p. 4842–4847.

Klimesch, W., 2012. Alpha-band oscillations, attention, and controlled access to stored information. Trends in Cognitive Sciences, 12, Volume 16, pp. 606–617.

Kluger, D. S. & Gross, J., 2020. Respiration modulates oscillatory neural network activity at rest.

Li, S. & Laskin, J. J., 2006. Influences of ventilation on maximal isometric force of the finger flexors. Muscle & Nerve, Volume 34, p. 651–655.

Li, S. & Rymer, W. Z., 2011. Voluntary Breathing Influences Corticospinal Excitability of Nonrespiratory Finger Muscles. Journal of Neurophysiology, 2, Volume 105, p. 512–521.

Lo, S. & Andrews, S., 2015. To transform or not to transform: using generalized linear mixed models to analyse reaction time data. Front. Psychol., August.

Montoya, P., Schandry, R. & Müller, A., 1993. Heartbeat evoked potentials (HEP): topography and influence of cardiac awareness and focus of attention. Electroencephalography and Clinical Neurophysiology/Evoked Potentials Section, 5, Volume 88, p. 163–172.

Moreno-Martińez, F. J. & Montoro, P. R., 2012. An Ecological Alternative to Snodgrass & Vanderwart: 360 High Quality Colour Images with Norms for Seven Psycholinguistic Variables. PLoS ONE, 5, Volume 7, p. e37527.

Mumford, D., 1992. On the computational architecture of the neocortex. Biol. Cybern., June, p. 241–251.

Myllylä, T. et al., 2017. Multimodal brain imaging with magnetoencephalography: A method for measuring blood pressure and cardiorespiratory oscillations. Scientific Reports, 3.Volume 7.

Nakamura, N. H., Fukunaga, M. & Oku, Y., 2018. Respiratory modulation of cognitive performance during the retrieval process. PLOS ONE, 9, Volume 13, p. e0204021.

Negro, C. A. D., Funk, G. D. & Feldman, J. L., 2018. Breathing matters. Nature Reviews Neuroscience, 5, Volume 19, p. 351–367.

Obleser, J. et al., 2021. Circadian fluctuations in glucocorticoid level predict perceptual discrimination sensitivity. bioRxiv, March.

Ondobaka, S., Kilner, J. & Friston, K., 2017. The role of interoceptive inference in theory of mind. Brain and Cognition, Volume 112, pp. 64–68.

Oostenveld, R., Fries, P., Maris, E. & Schoffelen, J.-M., 2011. FieldTrip: Open Source Software for Advanced Analysis of MEG, EEG, and Invasive Electrophysiological Data. Computational Intelligence and Neuroscience, Volume 2011, p. 1–9.

Park, H.-D.et al., 2020. Breathing is coupled with voluntary action and the cortical readiness potential. Nature Communications, 2.Volume 11.

Perl, O. et al., 2019. Human non-olfactory cognition phase-locked with inhalation. Nature Human Behaviour, 3, Volume 3, p. 501–512.

Petzschner, F. H. et al., 2019. Focus of attention modulates the heartbeat evoked potential. NeuroImage, 2, Volume 186, p. 595–606.

Rasch, D., 1986. Gibbons, J. D.: Nonparametric Statistical Inference. Biometrical Journal, 28(8), p. 936– 936.

Rousselet, G. A., Pernet, C. R. & Wilcox, R. R., 2017. Beyond differences in means: robust graphical methods to compare two groups in neuroscience. European Journal of Neuroscience, 6, Volume 46, p. 1738–1748.

Schroeder, P. A., Schwippel, T., Wolz, I. & Svaldi, J., 2020. Meta-analysis of the effects of transcranial direct current stimulation on inhibitory control. Brain Stimulation, 5, Volume 13, pp. 1159–1167.

Short, S. M. et al., 2016. Respiration Gates Sensory Input Responses in the Mitral Cell Layer of the Olfactory Bulb. PLOS ONE, 12, Volume 11, p. e0168356.

Stüttgen, H. H. M. C., 2011. Computation of measures of effect size for neuroscience data sets. European Journal of Neuroscience, November, pp. 1887–1894.

Suchotzki, K. et al., 2017. Lying takes time: A meta-analysis on reaction time measures of deception. Psychol Bull, 4, Volume 143, pp. 428–453.

Szucs, D. & Ioannidis, J. P. A., 2017. Empirical assessment of published effect sizes and power in the recent cognitive neuroscience and psychology literature. PLoS Biol., March.

Thut, G., Nietzel, A., Brandt, S. A. & Pascual-Leone, A., 2006. α-Band Electroencephalographic Activity over Occipital Cortex Indexes Visuospatial Attention Bias and Predicts Visual Target Detection. Journal of Neuroscience, 9, Volume 26, pp. 9494–9502.

Tort, A. B. L., Brankačk, J. & Draguhn, A., 2018. Respiration-Entrained Brain Rhythms Are Global but Often Overlooked. Trends in Neurosciences, 4, Volume 41, p. 186–197.

Tort, A. B. L. et al., 2018. Parallel detection of theta and respiration-coupled oscillations throughout the mouse brain. Scientific Reports, 4.Volume 8.

Varga, S. & Heck, D. H., 2017. Rhythms of the body, rhythms of the brain: Respiration, neural oscillations, and embodied cognition. Consciousness and Cognition, 11, Volume 56, p. 77–90.

Vlemincx, E. et al., 2010. Sigh rate and respiratory variability during mental load and sustained attention. Psychophysiology, 12, Volume 48, p. 117–120.

Wagenmakers, E.-J. & Farrell, S., 2004. AIC model selection using Akaike weights. Psychonomic Bulletin & Review, 2, Volume 11, p. 192–196.

Wilcox, R., 2012. Introduction to Robust Estimation and Hypothesis Testing.

Zelano, C. et al., 2016. Nasal Respiration Entrains Human Limbic Oscillations and Modulates Cognitive Function. The Journal of Neuroscience, 12, Volume 36, p. 12448–12467.

